# Distinct Neural Signatures of Multimodal Resizing Illusions: Implications for Chronic Pain Treatment

**DOI:** 10.1101/2023.01.18.524558

**Authors:** Kirralise J. Hansford, Daniel H. Baker, Kirsten J. McKenzie, Catherine E. J. Preston

## Abstract

Illusory body resizing typically uses multisensory integration to change the perceived size of a body part. Previous studies associate such multisensory body illusions with frontal theta oscillations and parietal gamma oscillations for dis-integration and integration of multisensory signals, respectively. However, recent studies support illusory changes of embodiment from visual-only stimuli. Multisensory resizing illusions can also reduce chronic pain, potentially through modulation of cortical body representations. This preregistered study (N=48) investigated differences between multisensory visuo-tactile and uni-modal visual resizing illusions using EEG. We hypothesised (1) stronger illusion in multisensory compared to uni-modal, and uni-modal compared to asynchronous (dis-integration) conditions, (2) greater parietal gamma during multisensory compared to uni-modal, and (3) greater frontal theta during asynchronous compared to baseline conditions. Results partially supported EEG hypotheses, finding increased parietal gamma activity comparing multisensory to unimodal visual conditions, whilst finding increased parietal theta activity when comparing asynchronous to non-illusion conditions. While results demonstrated that only 27% of participants experienced the illusion with visual-only stimuli, further analysis suggested that those who experience visual-only illusions exhibit a different neural signature to those who do not. Our results support the importance of multisensory integration for illusory changes in perceived body size. However, we also suggest that visual-only illusions can influence cortical body representations for a significant proportion of participants, which may have implications for the development of accessible visual-only chronic pain treatments.

## 1. Introduction

Illusory resizing is a form of multisensory illusion, often using visual and tactile inputs, whereby a body part is resized using augmented reality or magnifying optics and can consist of stretching or shrinking manipulations. Resizing illusions change how a body part looks to try to induce changes to cortical representations and subjective embodiment of the newly sized body part. Such illusory manipulations of the bodily self stem from studies using the rubber hand illusion (RHI). The RHI involves delivering tactile stimulation to a seen fake hand at the same time and in the same place that tactile stimulation is given to the hidden real hand, which elicits feelings of ownership over the fake hand. The integration of the multisensory (tactile and visual) inputs drives this illusory experience and taps into the neural substrates of our sense of bodily self, highlighting its apparent malleability (Botvinick & Cohen, 1998). Leading from these findings, further research has shown that embodiment can also occur during mirror illusions, such as those used in phantom limb studies (Chan et al., 2007), in which a mirror is placed adjacent to the patient’s remaining limb, giving an illusion of the amputated limb still being there, and from multisensory resizing illusions involving both tactile and visual inputs within augmented reality manipulations (Preston and Newport, 2011).

In addition to multisensory resizing illusions, embodiment has also been reported for unimodal visual resizing illusions such as when viewing an illusion of an elongated arm (Schaefer et al., 2007), while changes to embodied perception have also been reported from visual-only manipulations of the hand (McKenzie & Newport, 2015) and illusory experience has been successfully induced in the rubber hand illusion using visual-only stimulation (Ferri et al., 2014). Furthermore, visual capture alone has been found to elicit embodiment in both immersive virtual (Maselli & Slater, 2013) and physical environments (Carey et al., 2019) for full body illusions. Interestingly, it has been found that some individuals are more susceptible than others to visual only manipulations, with some participants not experiencing embodiment at all (Carey et al., 2019). The subjective embodiment measures used in these studies have primarily consisted of self-report questionnaires, with limited research into the accompanying neural responses to such illusions.

Of the previous studies looking at EEG data and multimodal information processing, the parietal area specifically has been proposed as a multimodal integration processing site (Kanayama et al., 2007). This is due to studies demonstrating a relationship between gamma-band oscillations and integration of multisensory processes across both auditory and visual stimuli (Kaiser et al., 2005; Sakowitz et al., 2005; Senkowski et al., 2005). Looking specifically at visuotactile manipulations, such as those used in multisensory hand-based illusions (e.g., the rubber hand illusion), power increases have been observed in the gamma band (40–50 Hz) in parietal regions 200–250 ms into congruent visuotactile stimulation (Kanayama, 2007) in virtual and real-life environments (Kanayama et al., 2021). This is posited to reflect an early stage of multimodal stimulus integration, highlighting parietal regions as potential seats of multisensory integration. fMRI findings from Ehrsson et al. (2005) who delivered the rubber hand illusion in MRI scanners, and from Petkova et al. (2011), who used body-swap illusions in an fMRI study, also support parietal involvement in multisensory integration, finding activity in the ventral premotor cortices, intraparietal cortices, and the cerebellum (Ehrsson et al., 2005) in addition to the bilateral ventral premotor cortex, the left intraparietal cortices and the left putamen (Petkova et al., 2011; Preston & Ehrsson, 2016). Multisensory EEG research has also pointed to the existence of oscillatory components related to multisensory integration within theta bands. Theta band (3–8 Hz) activity has been observed between 100 and 300 ms post stimulus (Kanayama et al., 2021). Whilst gamma band activity is observed around the parietal region and shows greater activity to spatially congruent visuo-tactile tasks, theta band activity is found around frontal sites and shows greater response to spatially incongruent visuo-tactile stimulation. Increases in theta power have been attributed to the cognitive load required to process incongruent visuotactile information (Kanayama et al., 2021). Research further supporting the frontal location of theta activity comes from Petkova et al. (2011), who used a full body ownership illusion and fMRI, and found increased activity in the ventral premotor cortex linked to construction of ownership of the body, cognitive load, and control processes.

In addition to being a useful method to investigate the malleability of our bodily self, resizing illusions have also shown the potential to reduce pain in chronic pain conditions such as complex regional pain syndrome (CRPS) (Moseley, Parsons & Spence, 2008), chronic back pain (Diers et al., 2013) and osteoarthritis (OA) of the hand (Preston & Newport, 2011) and knee (Stanton et al., 2018). Theories regarding this pain reduction are linked to the inaccurate size reports chronic pain patients often give to their affected limbs (Lewis et al., 2007; Moseley, 2005; Peltz et al., 2011; Gilpin et al., 2014; Stanton et al. 2018) and the resizing illusions ameliorating this discordance. Multisensory illusory resizing, however, requires the use of a large augmented-reality system as well as the presence of a researcher to deliver the manipulations, and is therefore, somewhat impractical as a treatment option. Given that unimodal visual illusions have also been shown to elicit subjective embodiment, it is plausible that there could be accompanying analgesia. Recently, there has been evidence to suggest that visual-only illusory resizing of the hand in complex regional pain syndrome can reduce pain levels (Lewis et al., 2021), however, the neural underpinnings of both multisensory and unimodal visual resizing illusions are not yet understood, meaning that inferences regarding possible analgesic effects of unimodal visual resizing illusions in chronic pain more widely cannot be made.

Consequently, this study aims to further develop our understanding of the neural underpinnings of multimodal integration in a healthy participant sample by using EEG, in addition to subjective experience questionnaires, to enhance our understanding of the mechanisms behind resizing illusions in healthy participants and provide a foundation for future work with chronic pain patients. This will be achieved by investigating the neural signatures of multisensory and unimodal resizing illusions to determine whether the multi-modal aspects of the finger stretching / shrinking illusion used in previous augmented-reality illusions, notably the touch and the visual manipulation of hand / finger size, are required for induction of the illusory experience, or if a uni-modal visual-only illusion is also able to elicit similar levels of illusory experience. If similar levels of illusory experience can be elicited by a uni-modal visual manipulation, it suggests that the analgesia experienced during visuotactile illusions could also be experienced during visual-only illusions. Given the previous literature denoting the feasibility of uni-modal visual illusions, the first hypothesis for the study is that (i) illusion strength will be greater in the multi-sensory (MS) condition compared to the uni-modal visual (UV) condition, which will be greater than an asynchronous control (AS) condition. Referring to the neural underpinnings of these illusions, the next hypothesis is that (ii) there will be stronger parietal gamma band activity (30 – 60Hz) elicited during MS compared to UV conditions, and finally, to assess additional cognitive demands of the asynchronous condition, (iii) there will be greater frontal theta activity (5 – 7Hz) elicited during AS conditions compare to a non-illusion baseline condition.

## 2. Methods

### 2.1 Data / Code Availability Statement

Raw EEG data for each participant and the code used to analyse the data can be found at the following OSF link: https://osf.io/7wpqe/ DOI:10.17605/OSF.IO/7WPQE

### 2.2 Ethics Statement

Ethical approval for this study was granted by the Ethics Committee of the Department of Psychology at the University of York. All participants gave written informed consent before taking part.

### 2.3 Participant Sample

#### 2.3.1 Power Analysis and Sample Size

A priori power analysis using illusion data from a pilot study showed a minimum sample size of 26 participants was required (d = .67, power = .95, alpha = .05). Due to the small pilot study sample size (n = 9) and the current study using EEG, which was not used previously, in addition to the inherent ambiguity of power analyses and to account for participant drop out / attrition, the sample size of 26 participants was approximately doubled, with recruitment of 50 participants.

#### 2.3.2 Participants

48 participants (83.5% Female. 14.5% Male, 2% Non-Binary; Mean age = 21 years) completed the experiment (2 participants lost to drop out / attrition), with exclusion criteria being prior knowledge or expectations about the research, a history of neurological or psychiatric disorders, operations or procedures that could damage peripheral nerve pathways in the hands, a history of chronic pain conditions, history of drug or alcohol abuse, history of sleep disorders, history of epilepsy, having visual abnormalities that cannot be corrected optically (i.e. with glasses), or being under 18 years of age.

### 2.4 Materials

Participants were fitted with a 64-channel EEG cap (ANT Neuro Waveguard) with electrodes arranged according to the 10/20 system. EEG set up included use of conductive gel between the electrodes and the scalp to attempt to obtain impedance levels of <10kΩ per electrode. Resizing illusions were delivered using an augmented-reality system (see Figure 1) that consisted of an area for the hand to be placed which contained a black felt base, LED lights mounted on either side and a 1920 × 1080 camera situated in the middle of the area, away from the participant’s view. Above this area, there was a mirror placed below a 1920 × 1200 resolution screen, so that the footage from the camera was reflected by the mirror such that the participant could view live footage of their occluded hand. The manipulation of the live feed from the camera was implemented using MATLAB r2017a, wherein the participant’s finger would stretch / shrink by 60 pixels during illusions lasting 2.4 seconds. This stretching or shrinking would be accompanied during the multisensory condition by the experimenter gently pushing or pulling on the participant’s finger to induce immersive multisensory illusions. After manipulation, there was a 2.4 second habituation phase in which participants could view and move their augmented finger before the screen went dark, indicating that the next trial could start. Subjective illusion experience was collected via Qualtrics (Qualtrics, Provo, UT) on a Samsung Galaxy Tab A6 tablet. This was given to participants at towards end of the experiment, wherein each trial was presented again, and subsequently participants were asked to recall the trial they had just experienced and previous trials that were similar, and then give a response on a Likert scale of -3 to +3, with -3 being strongly disagree and +3 being strongly agree with statements made. The questionnaire consisted of six statements, two relating to illusory experience: “It felt like my finger was really stretching” / “It felt like the hand I saw was part of my body”, two relating to disownership: “It felt like the hand I saw no longer belonged to me” / “It felt like the hand I saw was no longer part of my body”, and two were control statements: “It felt as if my hand had disappeared” / “It felt as if I might have had more than one right hand”. The questionnaire was delivered 7 times, once after each trial.

**Figure 1.**
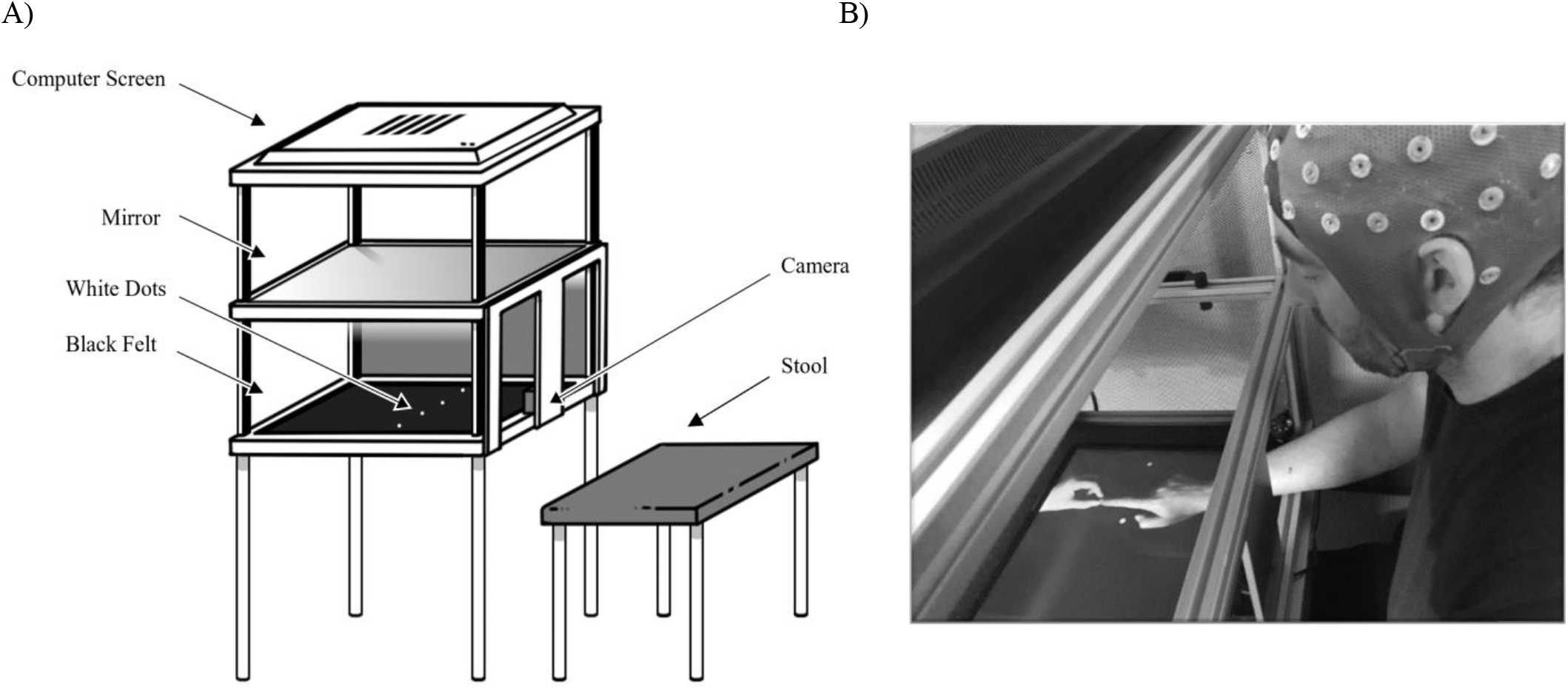
A) Schematic of Augmented Reality System. B) Image of Participant in EEG Cap Undergoing Resizing Illusion.

### 2.5 Procedure

After EEG set up, participants were seated at the augmented-reality system and instructed to place their right hand, with their index finger outstretched, onto the felt. There were two white dots on the felt to guide where their hand should be placed. Participants were instructed to view their hand’s image in the mirror (whilst their real hand was hidden from view) throughout the experiment. Participants completed 12 repetitions of 7 distinct conditions: 1, immersive multisensory (MS) stretching; 2, immersive multisensory (MS) shrinking; 3, unimodal visual (UV) stretching; 4, unimodal visual (UV) shrinking; 5, asynchronous control (AS) stretching; 6, asynchronous control (AS) shrinking; 7, non-illusion baseline. Multisensory conditions consisted of the experimenter pulling or pushing the participant’s index finger as the participant viewed their hand stretching or shrinking in a congruent manner. Unimodal conditions consisted of the participants viewing their finger either stretch or shrink without any experimenter manipulation. Asynchronous conditions consisted of the experimenter pushing or pulling the participant’s index finger as the participant viewed their hand stretching or shrinking in an incongruent manner. Non-illusion conditions provided no visual or tactile manipulations of the finger. (Video of a participant undergoing multisensory stretching can be seen in supplementary material). Conditions were randomised via MATLAB r2017a, and the experimenter was unaware which condition would be presented on a given trial. The experimenter was then informed of whether to push or pull the finger or to apply no manipulation via audio cues delivered through Bluetooth earphones. 6 repetitions of the 7 conditions were presented, followed by a break for the participant to stretch their hand and rest, and then another 6 repetitions of the 7 conditions were presented, again in a random order. There was then another break before each condition was presented once in a fixed order, after which the participant completed the subjective illusory experience questionnaire. EEG was recorded throughout as a continuous recording with conditions indicated by numbered triggers sent when the researcher pressed a button box to start the illusion for each trial.

### 2.6 Data Processing

#### 2.6.1 EEG Data Collection

EEG data were recorded continuously at 1kHz using the ASALab software. 8-bit digital triggers indicating trial onset and the end of the habituation period were sent from the stimulus computer to the EEG amplifier using a USB TTL module (Black Box Toolkit Ltd., UK).

#### 2.6.2 Questionnaire Data Collection

A Samsung Galaxy Tab A6 tablet was used to collect subjective illusory experience data via a questionnaire on Qualtrics (Qualtrics, Provo, UT), which the participants completed themselves, with a researcher present to answer any questions.

### 2.7 Data Analysis

#### 2.7.1 EEG Data Analysis

To identify noisy data, we calculated the standard error over time (Luck et al., 2021) for each electrode for each participant (following application of a 50Hz notch filter). Any electrode with a standard error in the top 5% of values (here, above a standard error of 1.5µV), or with a value of 0, were removed from the main analysis. Where a participant had over 50% of their electrodes over the 1.5 standard error threshold, their data were removed, resulting in a final sample of 47 participants (1 removed). The main EEG analysis was then conducted using Brainstorm (Tadel et al., 2011) where again a 50Hz notch filter was applied to the data, and trials were epoched to 5 seconds at intervals of 1000ms. Time-frequency analysis (Morlet wavelets) across trials was completed for each condition for each participant, with central frequency at 1Hz and time resolution (FWHM) at 3s. Data were grouped in frequency bands with the following ranges: Delta (2-4Hz), Theta (5-7Hz), Alpha (8-12Hz), Beta (15-29Hz), Gamma (30-60Hz). Arithmetic averages were then computed for each condition across all participants, and then again over both MS conditions, both UV conditions and both AS conditions. A pre-stimulus baseline period of 1000ms was included, and activity here was subtracted from all subsequent timepoints, leaving 5 experimental timepoints: 0 – 0999ms, 1000 – 1999ms, 2000 – 2999ms, 3000 – 3999ms, and 4000 – 5000ms. Changes in magnitude were statistically assessed using non-parametric cluster-based permutation analysis (Maris & Oostenveld, 2007) implemented in MATLAB r2017a. Here, a one-sample T test statistic and p-value were calculated for each sensor / time point, using a threshold of p < .05, before a list of clusters with significant elements was produced. The largest cluster was then stored, and a null distribution was built from 1000 random sets of permutations of the group condition labels and signs. The clusters were then compared to the null distribution and any clusters falling outside of the 95% confidence intervals were retained. The electrode within the significant cluster with the greatest effect size was then used to plot activity over the time course of the experiment, to illustrate the effect seen.

#### 2.7.2 Questionnaire Data Analysis

Raw data was exported from Qualtrics, and statistical analysis was completed in JASP (JASP Team, 2022). Scores for both illusion experience questions were averaged, along with both disownership questions and both control questions, resulting in 3 scores per trial per participant. Both MS conditions were then averaged, along with both AS and UV conditions, resulting in each participant giving 4 data points, one for MS, one for AS, one for UV and one for Baseline. Due to the nature of the Likert scale data not being continuous, a Friedman test was run to compare mean scores from each condition. Given significant findings, post-hoc Conover’s tests were run, with Bonferroni correction for 6 comparisons at an initial alpha of 0.05.

## 3 Results

Hypothesis 1 predicts that reported illusion strength will be greater in the MS condition compared to the UV condition, which will be greater than an AS condition. To test this, a Friedman test was conducted to determine whether illusion strength differed between MS, AS, UV and baseline conditions. Results, summarised in Figure 2A, show a significant difference between conditions (χ^2^(3) = 40.936, p <.001; W = .29), with posthoc Conover tests showing significant comparisons after Bonferroni correction between baseline and MS conditions (T(138) = 5.10, p <.001), MS and AS conditions (T(138) = 3.38, p = .006), and MS and UV conditions (T(138) = 5.86, p <.001). However, note that the illusion strength was *lower* in the UV condition than the AS condition, opposite to our hypothesis.

**Figure 2.**
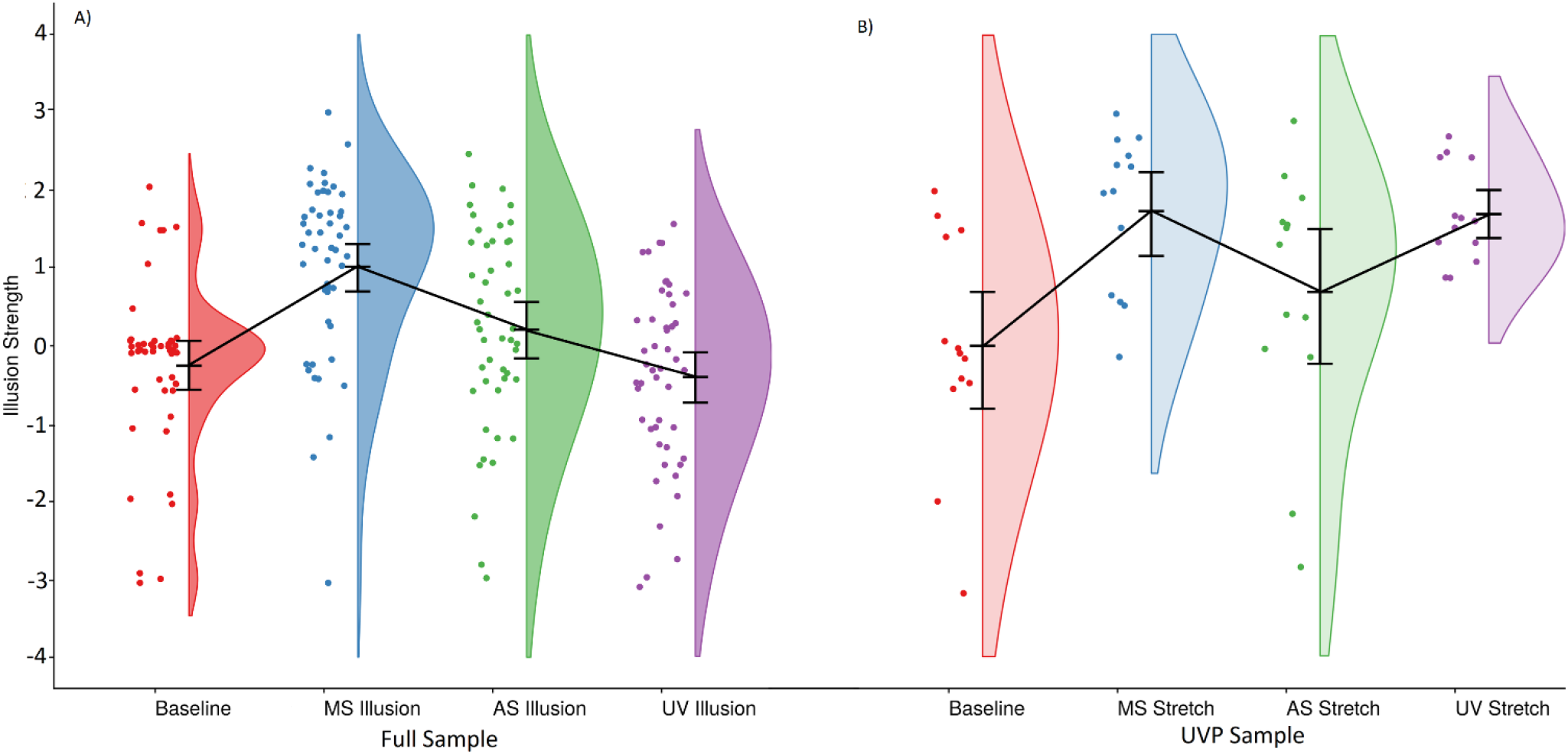
A) Illusion Strength in Each Averaged Illusion Condition for the Full Sample. B) Illusion Strength in Each Stretching Illusion Condition for the UVP Sample (27% of Participants). Error bars indicate 95% confidence intervals.

An average rating of ≥ +1 shows that the participants had an experience of ownership (this criterion has been used previously: Ehrsson et al., 2004; Petkova & Ehrsson, 2009; Kalckert & Ehrsson, 2012). Of the total participants, 13 out of 48 participants (27%) scored ≥ +1 on combined stretching illusion scores, showing an experience of the illusion in the UV stretching condition, and 5 out of 48 participants (10%) scored ≥ +1 on combined shrinking illusion scores, showing an experience of the illusion in the UV Shrinking condition. Therefore, to assess differences in illusion strength when there is an effective illustration of the UV condition, a Friedman test was conducted on the 27% of participants who experienced an effective UV stretching condition, now termed the uni-modal visual positive (UVP) sample, to determine whether illusion strength differed between MS, AS, UV and baseline stretching conditions. This exploratory analysis was not, however, conducted on the 10% who experienced an illusion in the UV shrinking condition, since the sample size was so small that power to detect meaningful effects was minimal.

Results, summarised in Figure 2B, show a significant difference between conditions (χ^2^(3) = 13.703, p =.003; W = .351), with post-hoc Conover tests showing significant comparisons after Bonferroni correction between baseline and MS Stretch (T(36) = 3.40, p = .002), baseline and UV Stretch (T(36) = 2.61, p = .013), and MS Stretch and AS Stretch (T(36) = 2.14, p = .039). Note that the illusion strength was *higher* in the UV condition than the AS condition in this group, in line with our hypothesis.

We next assessed Hypothesis 2, that there will be stronger parietal gamma band activity elicited during MS compared to UV conditions. A significant cluster comparing these conditions was found in the gamma band (30-60Hz) between 4000 and 5000ms (*p* = .008). The effect was strongest at electrode TP7, consistent with our prediction of a difference in parietal activity.

Due to the UV condition being present in this analysis, an exploratory analysis using the 27% of the sample who experienced an effective UV condition was also conducted. Here, three significant clusters were found in the gamma band between 0 and 1000ms (*p* < .001; *p* = .015; *p* < .001), again for comparing the MS and UV conditions. The difference was greatest over electrode F1, with clusters located in both frontal and parietal regions (see Figure 4).

Finally, hypothesis 3 predicted that there would be greater frontal theta (5-7Hz) activity elicited during AS conditions compared to a non-illusion baseline condition. A significant cluster was found in the theta band between 0 and 1000ms (*p =*.005) when comparing these two conditions. The difference was greatest over electrode M2, opposing our location prediction.

## 4 Discussion

This study aimed to further develop our understanding of the neural underpinnings of multimodal integration in healthy participants by using EEG in addition to subjective experience questionnaires across multisensory visuotactile, uni-modal visual, asynchronous, and non-illusion conditions. Findings demonstrated that reported illusion strength of the newly resized finger was found to be significantly stronger in multisensory compared to asynchronous and uni-modal visual conditions, and exploratory analysis highlighted that when there was an effective experience of the uni-modal condition, respective subjective embodiment surpassed that of the asynchronous condition. EEG analysis found increased gamma band activity in multisensory visuotactile compared to uni-modal visual conditions, in line with previous findings. This increased gamma therefore likely reflects multimodal stimulus integration effects, as the multimodal condition included visual and tactile manipulations, whereas the unimodal visual only included visual manipulations. Increased theta band activity was observed in the asynchronous compared to the non-illusion condition, likely reflecting additional cognitive load requirements to integrate conflicting sensory inputs. This increase in theta band activity was located in the parietal region, contrasting previous findings of frontal theta activity in asynchronous conditions.

Illusory experience data, as seen in Figure 2a, show a significant difference between conditions with a medium effect size (Lovakov & Agadullina, 2021), with an increase in subjective illusory experience in multisensory compared to unimodal conditions, as expected. Surprisingly, however, the asynchronous condition induced a stronger illusion than the uni-modal condition, in contrast to our first hypothesis. This unexpected finding can be explained through two possible ideas. First, the asynchronous condition might not have acted as an effective incongruent manipulation. This could be because during asynchronous stretching, where the participant’s finger stretched whilst the experimenter gently pushed the finger, this could instead act as a congruent multisensory condition, as the feeling of the experimenter pushing on the finger could feel as though the finger is pushing through a barrier, still giving a congruent stimulation effect. Exploratory analysis on disaggregated asynchronous data supports this idea, as there was a significant difference between the asynchronous stretching and asynchronous shrinking conditions (χ^2^(1) = 5.444, p = .02; W = .113), with post-hoc Conover tests showing significant comparisons after Bonferroni correction between AS Stretch and AS Shrink (p = .023), with participants experiencing a mean illusion strength score of .40 in AS Stretch compared to a mean illusion strength score of .01 in AS Shrink. Participants would be expected to show illusion scores of around 0, showing no illusory experience for an effective demonstration of the asynchronous condition. Therefore, the AS stretch condition (where the finger stretches visually but is compressed haptically) is likely to be a less appropriate control manipulation than the AS shrink condition (where the finger shrinks, but is stretched haptically). Secondly, as can be seen in Figure 2b, when participants do experience an effective uni-modal visual condition, as was the case with almost a third of our participants, the data show trends towards supporting our first hypothesis-that illusion strength would be greater in MS compared to UV, which would be greater than AS, with a slightly greater effect size than the full sample analysis. This, therefore, shows individual difference effects in the susceptibility of participants to experience a uni-modal visual illusion. Previous research has also found similar effects for visual only observation of a mannequin body, showing that 40% of participants experience subjective embodiment from visual-only observations (Carey et al., 2019). Furthermore, McKenzie and Newport (2015) found variability in the degree to which people experienced visually-induced sensations, finding a correlation between somatoform dissociation and visually-induced sensations. These findings suggest that if resizing illusions are to be used as an analgesic treatment option for chronic pain populations, it may be important to measure a patient’s susceptibility to uni-modal visual illusions before offering any kind of therapeutic intervention.

EEG data regarding multisensory integration can be seen in Figures 3 and 4. Findings show that for the total sample, a significant cluster is observed in the gamma band (30-60Hz) within the final phase of the experiment, which extends to parietal regions and possibly indicates a late stage of multimodal stimulus integration, expanding on previous findings regarding earlier stages of integration (Kanayama, 2007; Kanayama et al., 2021). It is possible that we see differences in the temporal nature of multimodal stimulus integration for a few reasons. Firstly, this study used illusory finger resizing as a method of multisensory manipulation, whereas previous studies have used the rubber hand illusion (Kanayama et al., 2021; Kanayama et al., 2007; Kanayama, Sato & Ohira, 2009; Hiramoto et al., 2017), or visual and tactile discrimination tasks (Kanayama & Ohira, 2009). Therefore, the differences seen in the gamma-band concerning early and late-stage multimodal stimulus integration could be due to different aspects of multisensory integration that are indexed by these different multisensory manipulations. Specifically, the integration in the rubber hand illusion differs from the integration in resizing illusions, as the rubber hand illusion elicits congruent tactile stimulation from the start of manipulation, whereas the resizing illusions elicit congruent tactile stimulation for a longer period as the finger resizes. Furthermore, in the present study, the gamma-band was classified as between 30 – 60Hz, whilst the previous studies which have observed significant increases in early-stage gamma-band power, have done so in more specific frequency ranges of 40-50Hz (Kanayama, 2007), 40-60Hz (Kanayama & Ohira, 2009) and 25-35Hz (Kanayama, Sato & Ohira, 2009). Additionally, Kanayama, Sato & Ohira (2009) found low-frequency gamma power reduction in congruent conditions, although non-significant, when dividing participants into groups based on depersonalisation tendencies. Hiramoto et al. (2017) suggested that reduction in low-frequency gamma could be modulated by individual differences. Our results also show individual differences in gamma activity, as seen in Figure 4 regarding the participant population who experienced an effective unimodal visual condition. Here, slightly decreased gamma activity was found in frontal regions, suggesting that those who experience visual-only resizing illusions demonstrate a different neural signature to those who do not. The significant clusters are in the manipulation phase, localised in both frontal and parietal regions. The difference in location of the significant clusters between the full sample and this subsample is likely due to the subsample experiencing an illusion in both multisensory and unimodal conditions, and therefore when looking at the difference in neural activity between the two conditions, this difference is seen at an early stage when there is the additional tactile input in the multisensory condition. Further analysis of the unimodal visual positive sample can be seen in supplementary materials.

**Figure 3.**
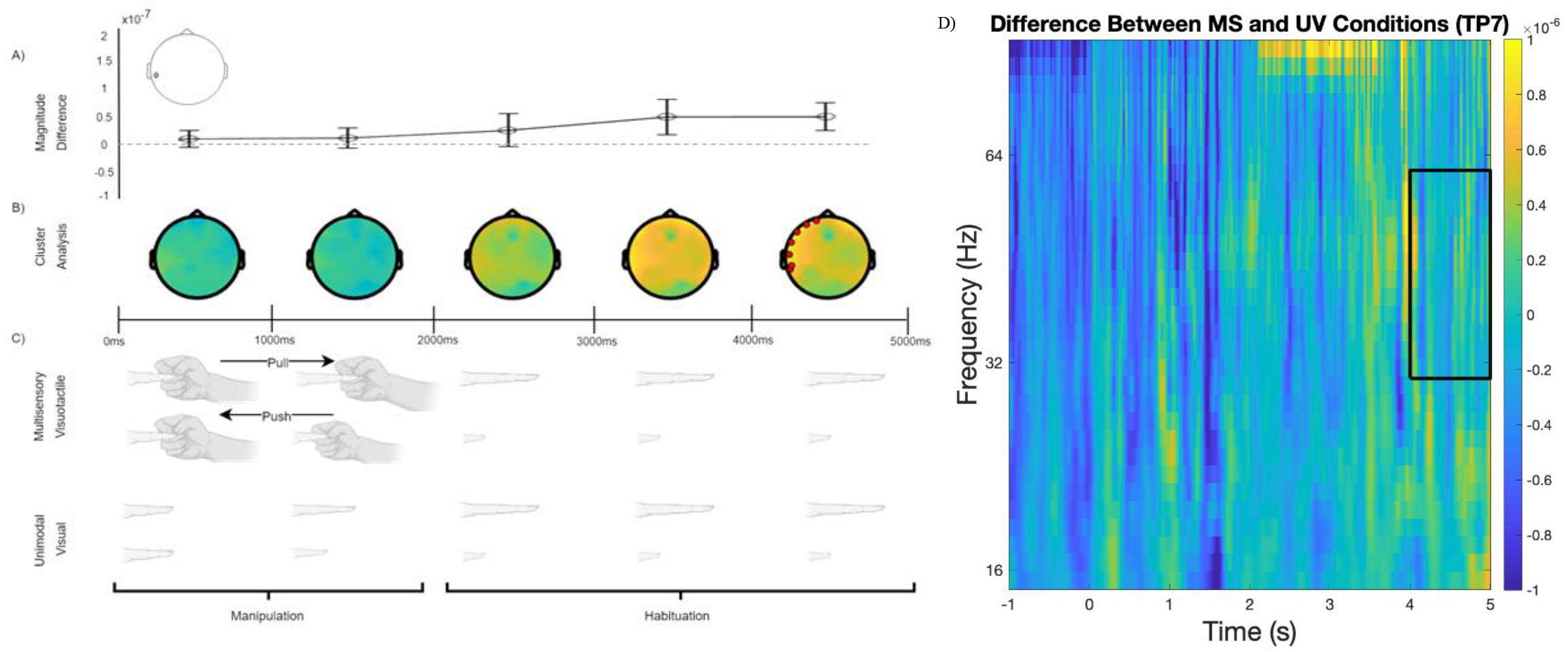
Comparison of gamma band activity between MS and UV conditions. The Magnitude Difference plot (a) shows time course of TP7 electrode, which was the significant electrode showing the largest effect size (d = 0.35). In panel (b), colour indicates the magnitude difference (blue: negative, yellow: positive), the significant clusters is highlighted by red dots. In panel (c), arrows denote the manipulation that the researcher’s hand is applying to the finger. Panel (d) shows the full time-frequency plot, with the black rectangle indicating the gamma band and the time-window containing the significant cluster.

**Figure 4.**
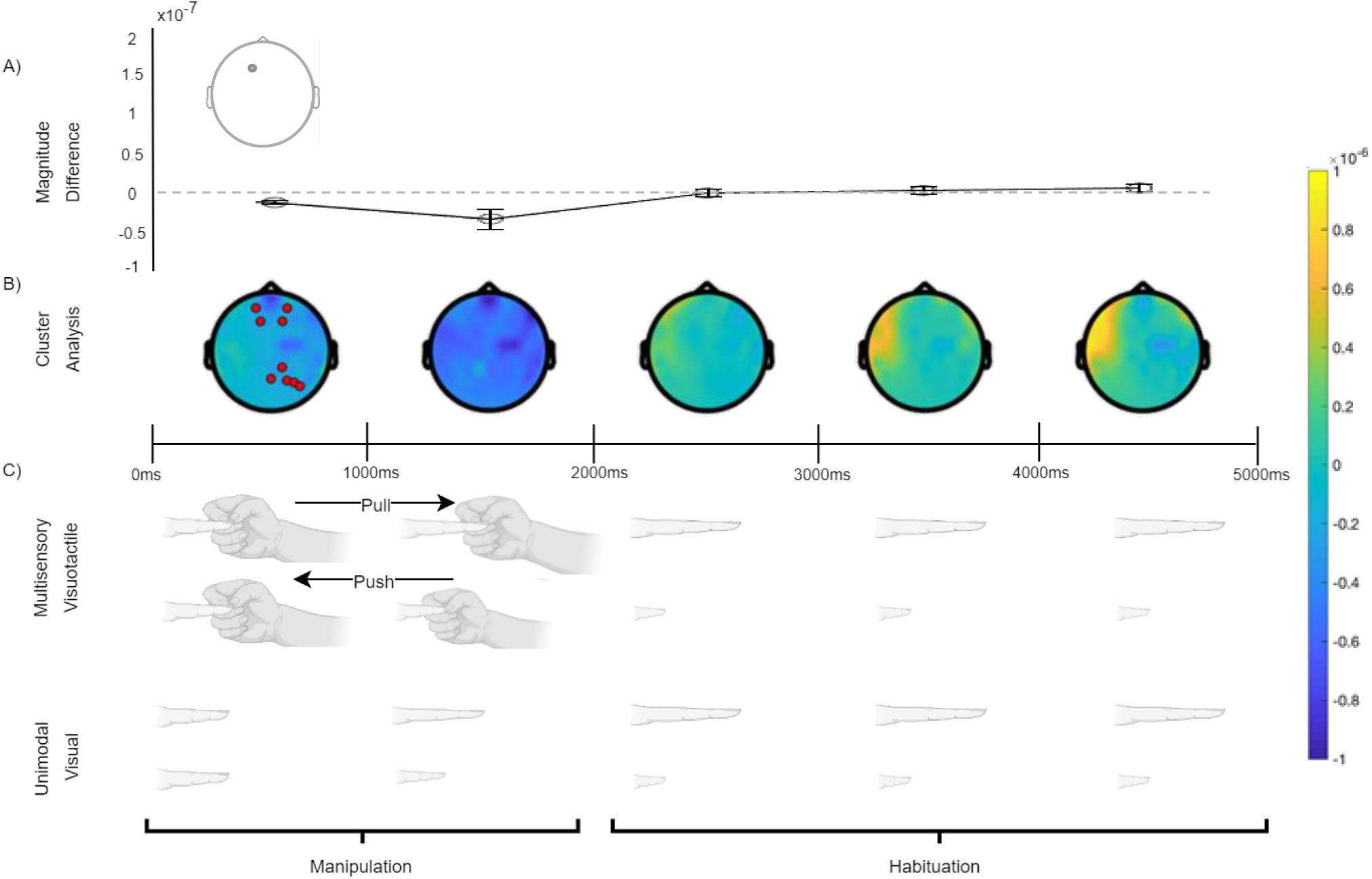
Comparison of gamma band activity between MS and UV conditions. The Magnitude Difference plot (a) shows time course of F1 electrode, which was the significant electrode showing the largest effect size (d = 0.91). In panel (b), colour indicates the magnitude difference (blue: negative, yellow: positive), and significant clusters are highlighted by red dots. In panel (c), arrows denote the manipulation that the researcher’s hand is applying to the finger.

EEG Data relating to multisensory dis-integration can be seen in Figure 5, which shows a significant cluster in the theta band (5-7Hz) 0 – 1000ms after onset of the manipulation. Previous literature posits that increases in theta band power relate to an additional cognitive load required to process the incongruent visuotactile information, which is likely reflected here in the theta band activity difference between asynchronous and non-illusion conditions., The increased theta band activity seen here is located around parietal sensors, ipsilateral to the tactile manipulation. This location contrasts with our hypothesis of increased frontal theta activity, however, this could be due to the aforementioned issue with the asynchronous stretching condition, whereby the finger is visually stretched whilst the researcher pushes on the finger, which could have been interpreted as the finger pushing against a barrier and therefore still feeling like a multisensory condition. This could then explain the parietal location, as multisensory integration effects have been previously linked to parietal areas (Kanayama et al., 2007). Additionally, EEG is known to lack discrete spatial resolution (Srinivasan, 1999), and therefore caution should be taken with this theta finding, and the previous gamma findings, when discussing the location of significant clusters.

**Figure 5.**
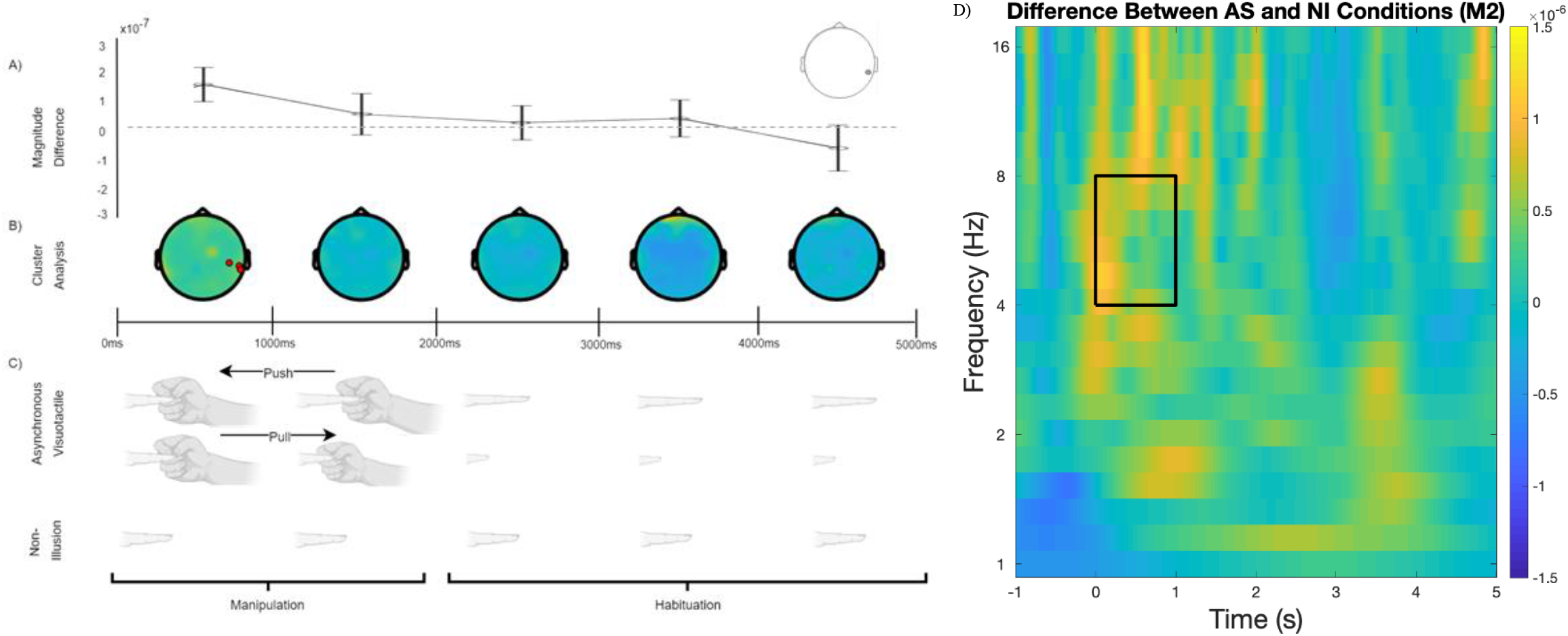
Comparison of theta band activity between AS and NI conditions. The Magnitude Difference plot (a) shows time course of M2 electrode, which was the significant electrode showing the largest effect size (d = 0.39). In panel (b), colour indicates the magnitude difference (blue: negative, yellow: positive), the significant cluster is highlighted by red dots. In panel (c), arrows denote the manipulation that the researcher’s hand is applying to the finger. Panel (d) shows the full time-frequency plot, with the black rectangle indicating the theta band and the time-window containing the significant cluster.

Taken together, our EEG findings support the previous literature regarding multisensory integration effects at gamma frequencies, and additional cognitive load requirements within the theta bands. Our findings enhance our understanding of the neural underpinnings of resizing illusions, showing that there could be important differences between multisensory visuotactile manipulations in rubber hand illusions and resizing illusions, relating to the temporal onset of integration effects. Our findings also add to the narrow previous literature regarding individual differences in gamma band power in multisensory conditions, showing here that a subset of participants who experienced an effective unimodal visual condition show spatially and temporally different effects compared to the full sample of participants, when comparing multisensory and unimodal visual conditions. These findings, however, could be enhanced by research investigating whether these illusions produce changes to the somatosensory cortex of participants. Neuroimaging has previously been used in healthy populations undergoing resizing illusions, wherein modulation of the primary somatosensory cortex has been found using neuromagnetic source imaging during resizing illusions of the arm (Schaefer et al., 2007). Given the differences seen between illusory resizing manipulations in these data, it is possible to posit that there will also be somatosensory cortex changes during finger resizing. There is also scope to investigate the differences between healthy and chronic pain participants, to see if the discordance reported for chronic pain conditions between real and perceived limb size would affect their somatosensory representations during illusory finger resizing.

These findings not only enhance our understanding of the neural signatures of multisensory visuotactile, uni-modal visual and asynchronous resizing illusions in healthy participants, but also provide a foundation to explore the neural signatures of resizing illusions in chronic pain populations. Further research is required to investigate whether the discordance in perception of limb size seen in chronic pain populations could result in different neural signatures to a healthy population. If found, this could indicate neural differences between the conditions that resizing illusions could help ameliorate, or conversely could show no differences between the populations, indicating a possible placebo analgesic effect of resizing illusions. Regarding future research with chronic pain populations, our data show that almost a third of healthy participants experience subjective embodiment in a visual-only illusion, which is supported by previous research (Carey et al., 2019), however, it is not known if a similar proportion of individuals experiencing an effective uni-modal visual condition would be seen in chronic pain populations, which therefore gives merit for future research into subjective embodiment during visual-only conditions for this population.

## 5. Conclusions

Overall, our findings support our EEG hypotheses in relation to activity increases in the gamma and theta bands, with both gamma and theta findings extending to parietal regions. These findings enhance our understanding of the neural signatures of visuotactile, visual only, and asynchronous illusory resizing manipulations in healthy participants, by adding novel evidence regarding what happens in a different presentation of a multisensory visuotactile illusion. Findings also show partial support for the subjective illusory experience hypothesis and illustrate the importance of individual differences in illusory experience of the uni-modal visual condition. This finding is arguably most significant when looking to future clinical applications of the uni-modal visual condition, as it highlights the value of susceptibility measures being included prior to analgesic interventions.

## Supporting information

Finger Stretch Video

## 6. Acknowledgements

The authors would like to acknowledge B. Quinn, for creation of the schematic in Figure 1a.

