## Supplementary Figures for "Distinct Neural Signatures of Multimodal Resizing Illusions: Implications for Chronic Pain Treatment"

**Supplementary Material**

Additional analysis on the UVP sample:

We assessed MS Stretch compared to NI in the UVP sample. Two significant clusters were found when comparing these conditions in the gamma band (30-60Hz) between 0 and 2000ms ($p$ = .01; P <.001). The effect was strongest at electrode CPZ.

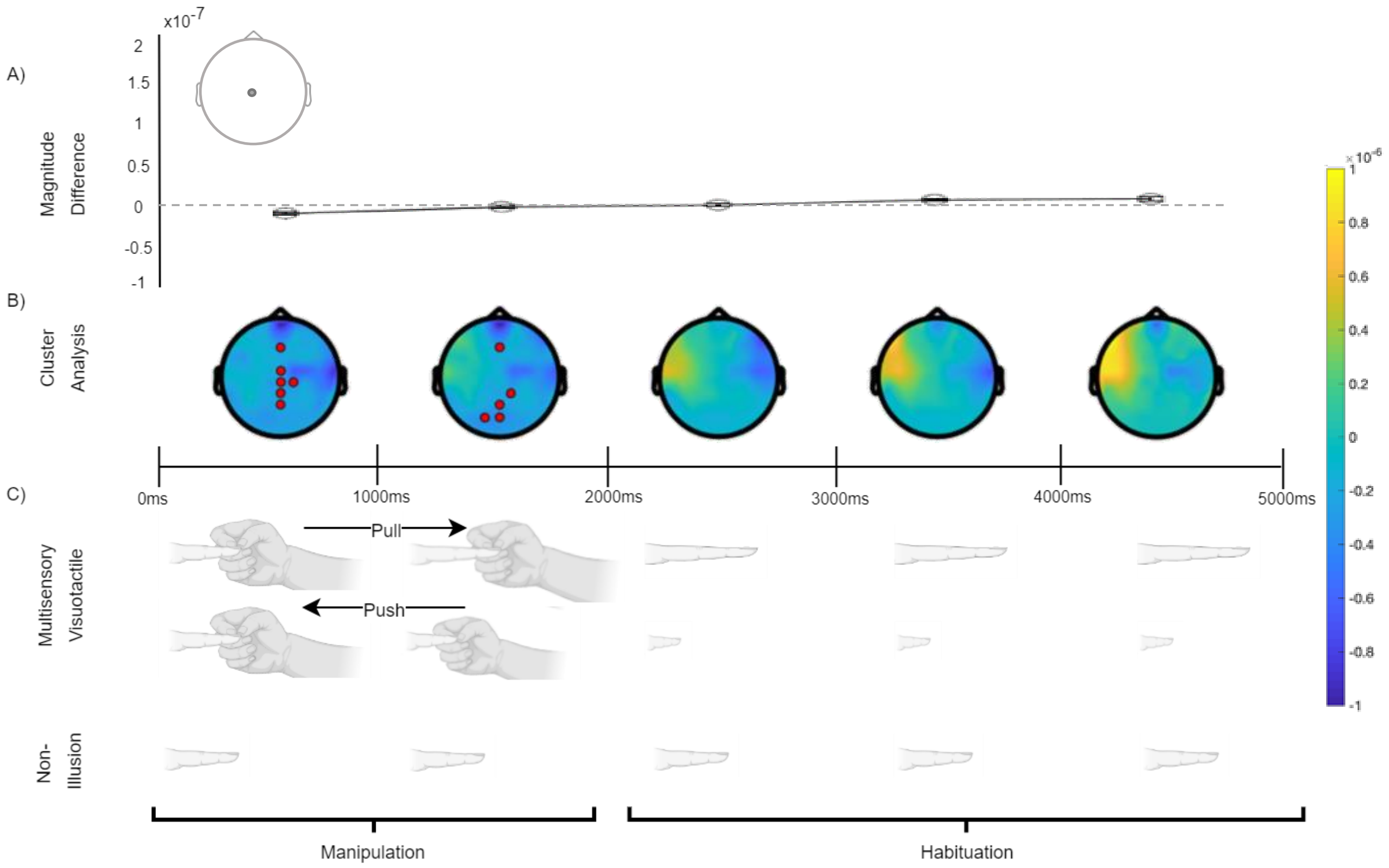


*Figure 7*. Comparison of gamma band activity between MS Stretch and NI conditions in the UVP Sample. The Magnitude Difference plot (a) shows time course of CPZ electrode, which was the significant electrode showing the largest effect size (d = 1.25). In panel (b), colour indicates the magnitude difference (blue: negative, yellow: positive), and significant clusters are highlighted by red dots. In panel (c), arrows denote the manipulation that the researcher's hand is applying to the finger.

We assessed UV Stretch compared to NI in the UVP sample. A significant cluster was found when comparing these conditions in the gamma band (30-60Hz) between 0 and 2000ms ($p$ = .012). The effect was strongest at electrode M1.

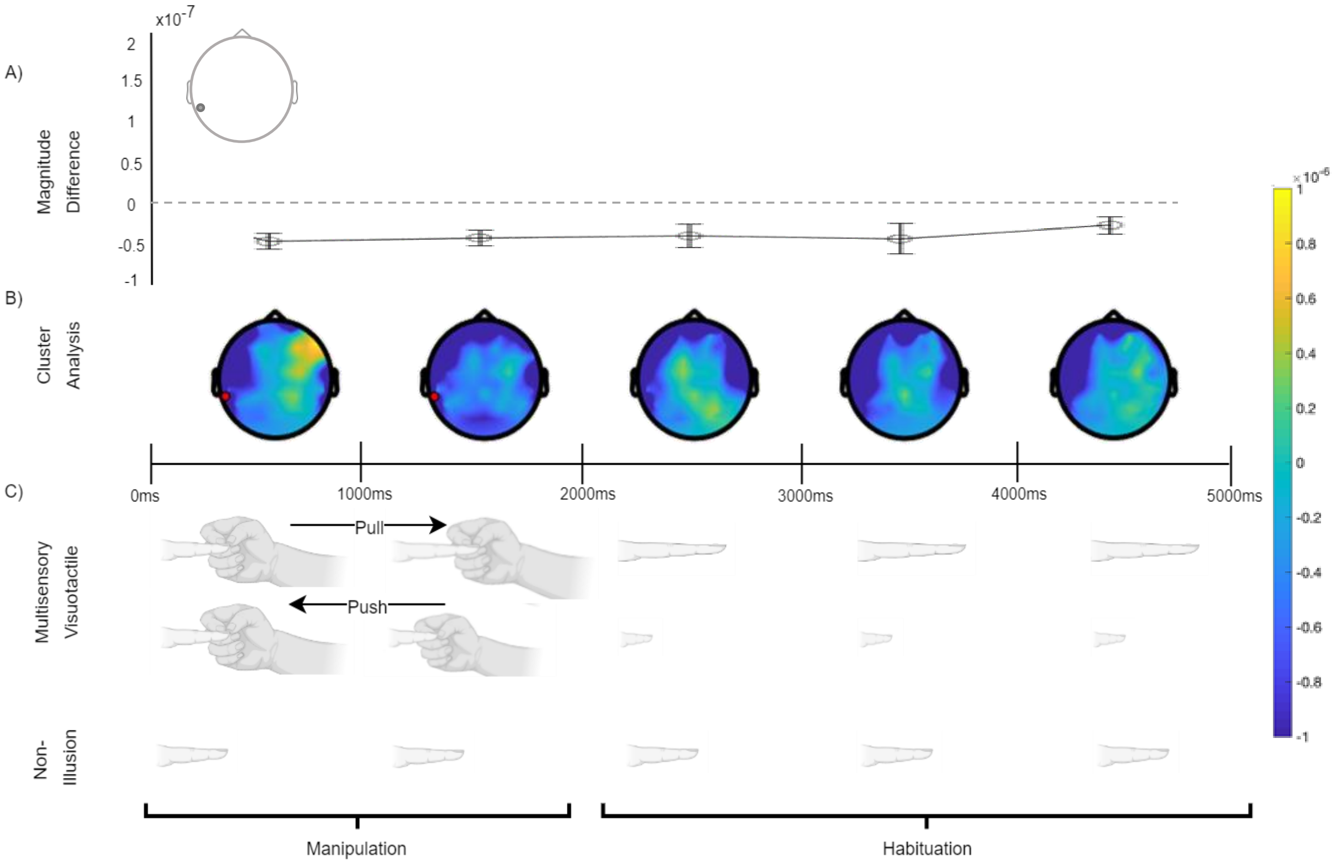


*Figure 8*. Comparison of gamma band activity between UV Stretch and NI conditions in the UVP Sample. The Magnitude Difference plot (a) shows time course of M1 electrode, which was the significant electrode showing the largest effect size (d = .98). In panel (b), colour indicates the magnitude difference (blue: negative, yellow: positive), and significant clusters are highlighted by red dots. In panel (c), arrows denote the manipulation that the researcher's hand is applying to the finger.

We assessed MS Shrink compared to NI in the UVP sample. Two significant clusters were found when comparing these conditions in the gamma band (30-60Hz) between 0 and 1000ms ($p$ = .002; P <.001). The effect was strongest at electrode F7.

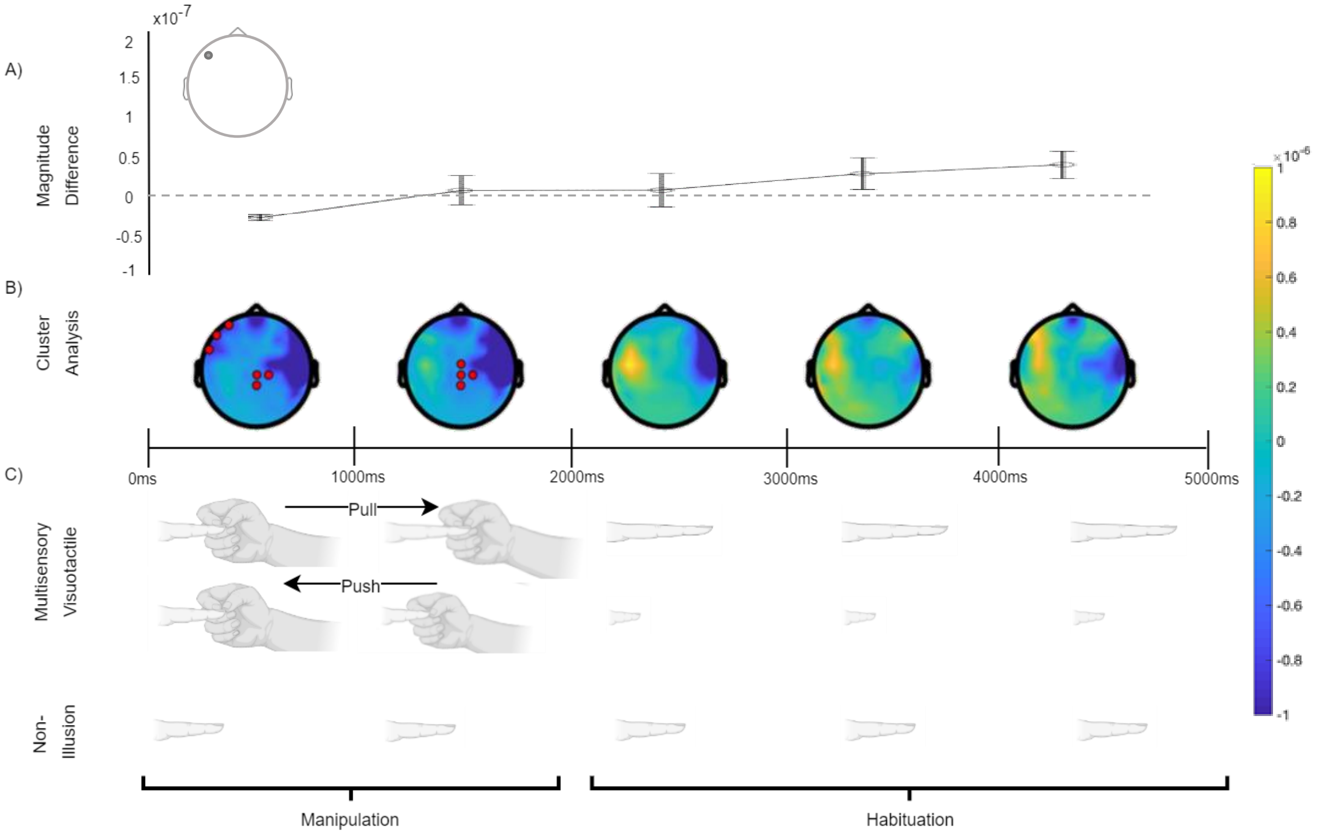


*Figure 9*. Comparison of gamma band activity between MS Shrink and NI conditions in the UVP Sample. The Magnitude Difference plot (a) shows time course of F7 electrode, which was the significant electrode showing the largest effect size (d = 1.15). In panel (b), colour indicates the magnitude difference (blue: negative, yellow: positive), and significant clusters are highlighted by red dots. In panel (c), arrows denote the manipulation that the researcher's hand is applying to the finger.

We assessed UV Shrink compared to NI in the UVP sample and found no significant clusters.
